# Translation elongation and mRNA stability are coupled through the ribosomal A-site

**DOI:** 10.1101/300467

**Authors:** Gavin Hanson, Najwa Alhusaini, Nathan Morris, Thomas Sweet, Jeff Coller

## Abstract

Messenger RNA (mRNA) degradation plays a critical role in regulating transcript levels in eukaryotic cells. Previous work by us and others has shown that codon identity exerts a powerful influence on mRNA stability. In *Saccharomyces cerevisiae*, studies using a handful of reporter mRNAs show that optimal codons increase translation elongation rate, which in turn increase mRNA stability. However, a direct link between elongation rate and mRNA stability has not been established across the entire yeast transcriptome. In addition, there is evidence from work in higher eukaryotes that amino acid identity influences mRNA stability, raising the question as to whether the impact of translation elongation on mRNA decay is at the level of tRNA decoding, amino acid incorporation, or some combination of each. To address these questions, we performed ribosome profiling of wildtype yeast. In good agreement with other studies, our data showed faster codon-specific elongation over optimal codons and faster transcript-level elongation correlating with transcript optimality. At both the codon-level and transcript-level, faster elongation correlated with increased mRNA stability. These findings were reinforced by showing increased translation efficiency and kinetics for a panel of 11 *HIS3* reporter mRNAs of increasing codon optimality. While we did observe that elongation measured by ribosome profiling is composed of both amino acid identity and synonymous codon effects, further analyses of these data establish that A-site tRNA decoding rather than other steps of translation elongation is driving mRNA decay in yeast.

## Introduction

The major eukaryotic mRNA degradation pathway is initiated by removal of the 3’ poly(A) tail (deadenylation), followed by cleavage of the 5’ 7mGpppN cap (decapping), and exonucleolytic degradation of the mRNA body in the 5’-3’ direction (Coller and Parker 2004; Ghosh and Jacobson 2010). Despite being targeted by a common degradation pathway, turnover rates for individual mRNAs differ dramatically, with halflives in yeast ranging from <1 minute to >60 minutes (Coller and Parker 2004). While RNA features in untranslated regions have been identified that influence the stability of some mRNAs (Muhlrad and Parker 1992; Lee and Lykke-Andersen 2013; Geisberg et al. 2014), we previously demonstrated that codon optimality (i.e., the balance between tRNA supply and codon demand) influences mRNA decay rates in a more global manner (Presnyak et al. 2015).

The tight coupling between translation status and mRNA turnover in dividing cells has long been appreciated (Jacobson and Peltz 1996; Coller and Parker 2004). Specifically, mRNA that is efficiently translated is more stable than mRNA that is translated poorly. The most parsimonious explanation for the link between translation efficiency and mRNA stability is that translation elongation rate is a major driver of mRNA decay rates. Indeed, *Saccharomyces cerevisiae* ribosome profiling studies have shown that cognate tRNA abundances correlate with translation efficiency (Hussmann et al. 2015; Weinberg et al. 2016), providing support that our observed correlation between codon optimality and mRNA stability may be due to differences in translation kinetics that feedback to the degradation machinery. Importantly, however, the links between codon optimality, translation efficiency, and mRNA stability have not been previously demonstrated on a genome-wide scale. In addition, since work in higher eukaryotes suggests that both A-site decoding and amino acid identity influence mRNA stability (Bazzini et al. 2016), we sought to determine which of these two aspects of translation elongation correlate more strongly with mRNA stability in *S*. *cerevisae*. In this study, we demonstrate that global codon- and transcript-level elongation rate estimates inferred by ribosome profiling correlate with mRNA stability. Further analysis of ribosome profiling and of translation efficiency and kinetics of reporter constructs indicate that A-site decoding links translation elongation to mRNA stability in yeast, while the amino acid effects on decay suggested in higher eukaryotes are either weaker or unrelated to translation elongation.

## Results

### Codon-specific translation elongation rates correlate with their influence on mRNA stability

In order to estimate relative codon-specific elongation rates, we generated a *Saccharomyces cerevisiae* ribosome profiling dataset using cells not pretreated with cycloheximide in order to accurately measure codon-level ribosome dynamics (Gerashchenko and Gladyshev 2014; Hussmann et al. 2015). As expected, a meta-analysis of ribosome protected fragments (RPFs) relative to the start of the coding sequence across all yeast mRNAs shows a clear periodicity of reads in frame with the coding sequence. The well-characterized pileup of reads 12 nucleotides upstream of the start codon is also present in 28 nucleotide RPFs (Fig. 1A). This is consistent with the start codon residing within the P-site of the ribosome, and we used this positioning to fix the location of the A-site within the reads (Ingolia et al. 2009). Relative per-codon ribosome elongation rates were estimated by utilizing a statistical framework that leverages linear mixed effects modeling of ribosome dynamics across a transcript (see Methods). This approach allows us to robustly model the error associated with ribosome density estimates and to quantify our confidence in the resulting estimates, while at the same time enabling us to relax arbitrary constraints on the data that we leverage to fit these models. Consistent with previous analyses (Hussmann et al. 2015; Weinberg et al. 2016), we find that our per-codon ribosome elongation rate estimates (ERE) correlate well (Pearson *r* = 0.55, *p* < 10^−5^) with the tRNA adaptation index (tAI), a measure of relative tRNA availability to the translation machinery of the cell (dos Reis et al. 2004) (Fig. 1B). This indicates that higher tRNA abundance is associated with more rapid elongation rates, presumably since the ribosome will more rapidly bind a codon’s cognate tRNA when that tRNA is more abundant within the cell.

**Figure 1.**
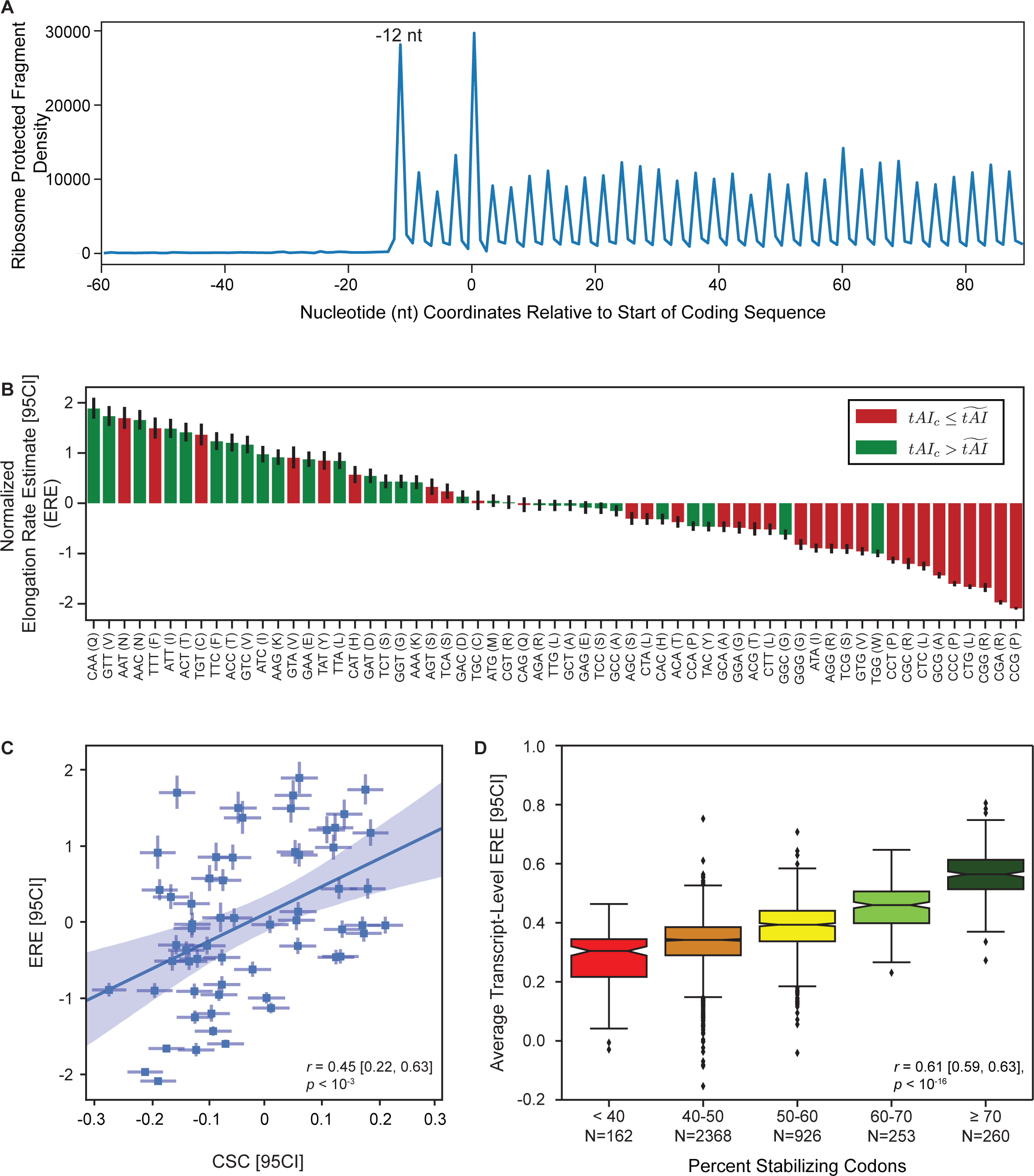
Codon-specific translation elongation rate estimates correlate with codon influence on mRNA decay. *(A)* Meta-gene analysis of 28 nucleotide ribosome protected fragments, relative to the first nucleotide of the coding sequence (position = 0). Read positions are counted based on the 5’ end of the ribosome protected fragments. *(B)* Normalized per-codon translation elongation estimates (EREs), ordered from fastest to slowest. The EREs for each of the 61 codons have been standardized to have a mean of 0 and a variance of 1. Error bars represent the 95% confidence intervals [95CI] for the estimate. We have colored each codon to reflect a codon-specific tAI value (tAI_c_) above (green) or below (red) the median tAI. *(C)* The best-fit line describing the relationship between normalized per-codon EREs and codon stability coefficients (CSCs), a measure of the influence of codons on mRNA stability. This relationship takes into account the uncertainty in the estimates of elongation rates and CSCs to arrive at a more robust estimate of the overall error in the correlation between these values. Uncertainties in both ERE and CSC values are included in the plot as the x- and y-axis 95% confidence intervals [95CI], respectively. The shaded region represents the 95% confidence intervals for the relationship between ERE and CSC. *(D)* Box plots showing the distribution of average transcript-level normalized EREs associated with the specified levels of percent stabilizing codons within mRNAs globally. Percent stabilizing codons reflects the proportion of codons in an mRNA with a CSC value greater than 0. Average transcript-level EREs are obtained by averaging per-codon EREs across the entire coding sequence of an mRNA. Notches reflect the standard error of the overall average EREs within each bin. Bin intervals are closed on the left and open on the right.

We next probed the relationship between our estimates of per-codon elongation rate and the codon occurrence to mRNA stability correlation coefficient (CSC), a measure of how individual codons contribute to the stability of mRNA transcripts (Presnyak et al. 2015). Consistent with the correlation previously observed between tAI and CSC values (Presnyak et al. 2015), we find a significant and positive relationship between ERE and CSC, both at the level of individual codons (*r* = 0.45 [0.22, 0.63], *p* < 10^−3^; Fig. 1C) and when elongation rates and CSC values are averaged across each mRNA (Pearson *r* = 0.61 [0.59, 0.63], *p* < 10^−16^; Fig. 1D). This latter analysis shows that mRNAs enriched in destabilizing codons are highly enriched in codons that facilitate slow ribosomal elongation.

Having shown a strong global relationship between ribosome elongation and the influence of codons on mRNA stability using our own data, we next sought to expand our analysis to examine this relationship using other ribosome profiling datasets. To this end, we analyzed ten *S*. *cerevisiae* ribosome profiling datasets generated without the use of cycloheximide (Cai and Futcher 2013; Gerashchenko and Gladyshev 2014; Guydosh and Green 2014; Jan et al. 2014; Lareau et al. 2014; Pop et al. 2014; Williams et al. 2014; Nedialkova and Leidel 2015; Young et al. 2015; Weinberg et al. 2016) (Supplemental Table S1) using the same analysis pipeline as our data. While the final normalized EREs varied across datasets, there is a clear partitioning of codons by elongation rate that is consistent across most of the datasets analyzed (Supplemental Table S1). A meta-analysis framework was adopted to estimate the true relationship between ERE and CSC values by aggregating the results obtained for each dataset into a population-level estimate of this relationship (Fig. 2A). This meta-analysis demonstrates that the body of generated ribosome profiling data in *S*. *cerevisiae*, considered in aggregate, is consistent with a significant relationship between per-codon elongation rate estimates and each codon’s influence on mRNA stability (r_Aggregate_ = 0.45 [0.34, 0.55], p < 10^−4^).

**Figure 2.**
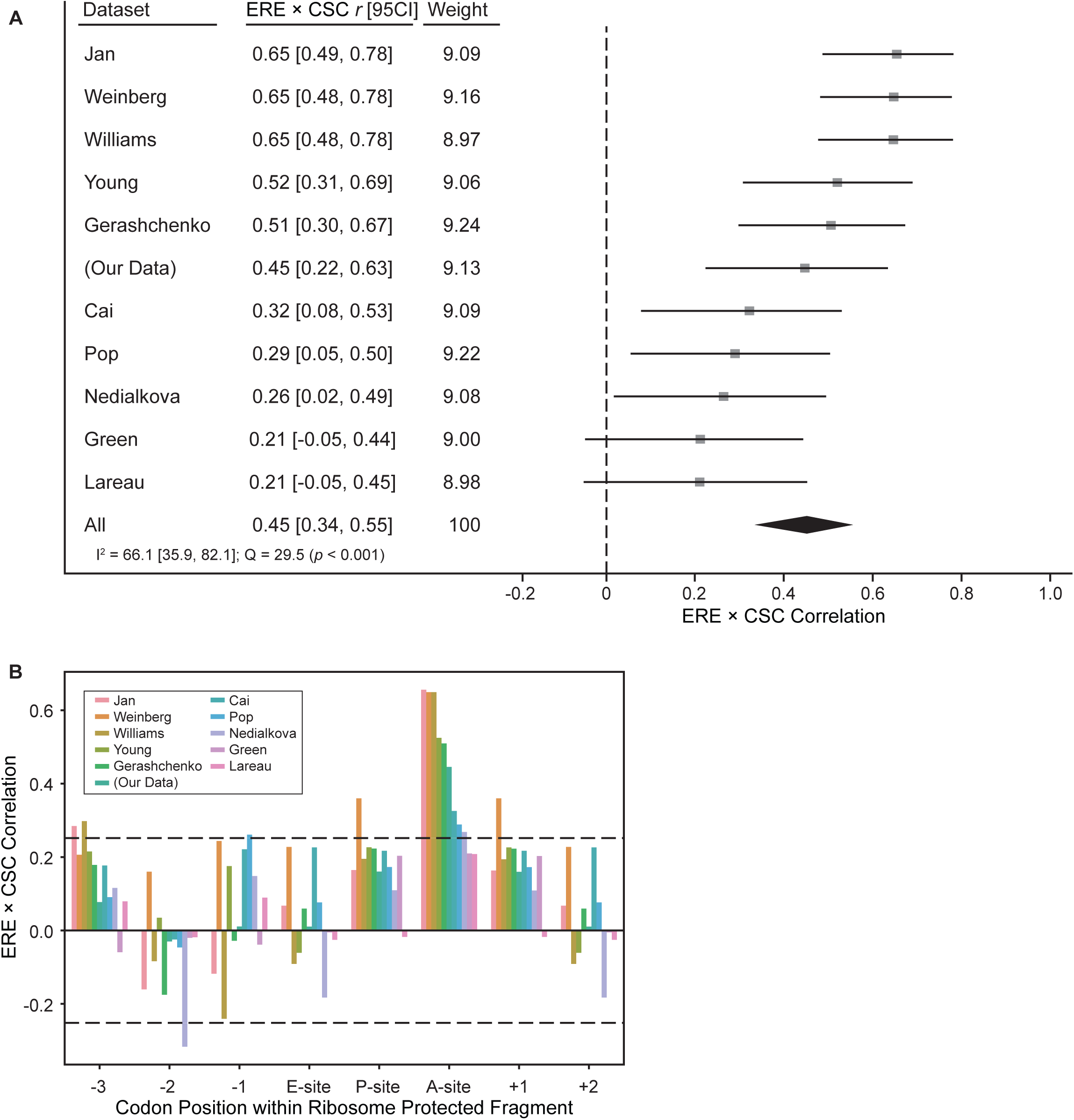
Translation elongation rates of codons within the ribosomal A-site correlate with codon influence on mRNA stability. *(A)* Forest plot describing the meta-analysis of the correlations between normalized elongation rate estimates (EREs) and codon stability coefficients (CSCs), a measure of the influence of codons on mRNA stability, using data from 11 cycloheximide-minus ribosome profiling experiments. The correlation between EREs and CSCs for each dataset are shown by the squares, with the error bars representing the associated 95% confidence interval [95CI]. The combined Pearson correlation estimate is represented by the large black diamond, with the width of the diamond representing the 95% confidence interval of the aggregate correlation estimate. I^2^ (the percentage of the variation across studies that is attributable to true between-study heterogeneity) and Cochran’s Q are also reported as standard measures of heterogeneity. *(B)* Bar plot showing the correlation between ERE and CSC values of codons located at specific locations within the ribosome protected fragments. The E-site, P-site, and A-site correspond to the trinucleotide sequence located at the +9, +12, and +15 positions within the ribosome protected fragments respectively, and this analysis is extended beyond these positions within the ribosome in either direction. The dashed lines represents the critical *r* value, such that values outside of the dashed lines represent statistically significant correlations at an uncorrected *α* = 0.05.

To test whether the observed codon effects are specific to codons located in the A-site of the ribosome, we repeated our analysis for a range of positions relative to the A-site, including the E-site and P-site, as well as regions further upstream and downstream of these sites (Fig. 2B). This allows us to test where the observed relationship between codon identity and transcript stability (CSC) occurs within the ribosome footprint. Only when codons are located in the A-site does the observed relationship between codon-specific EREs and CSCs correlate maximally, consistent with previous observations that codon identity within the A-site is uniquely associated with tAI (Weinberg et al. 2016).

### Ribosomal A-site decoding mediates the link between elongation rate and mRNA stability

The codon stability coefficient (CSC) is only an estimate of how codon content may be associated with transcript stability. More directly relevant is the relationship between the average elongation rate estimates across a transcript and that transcript’s stability, as determined by mRNA half-life measurements. Using published mRNA half-life data (Presnyak et al. 2015), we globally assessed the relationship between average ERE across a transcript and that transcript’s half-life. We find a significant and positive relationship between average elongation rate estimates and transcript half-life in our data (*r* = 0.28 [0.25, 0.31], *p* < 10^−16^; Fig. 3A) as well as in all datasets analyzed (Fig. 3B; *red*).

**Figure 3.**
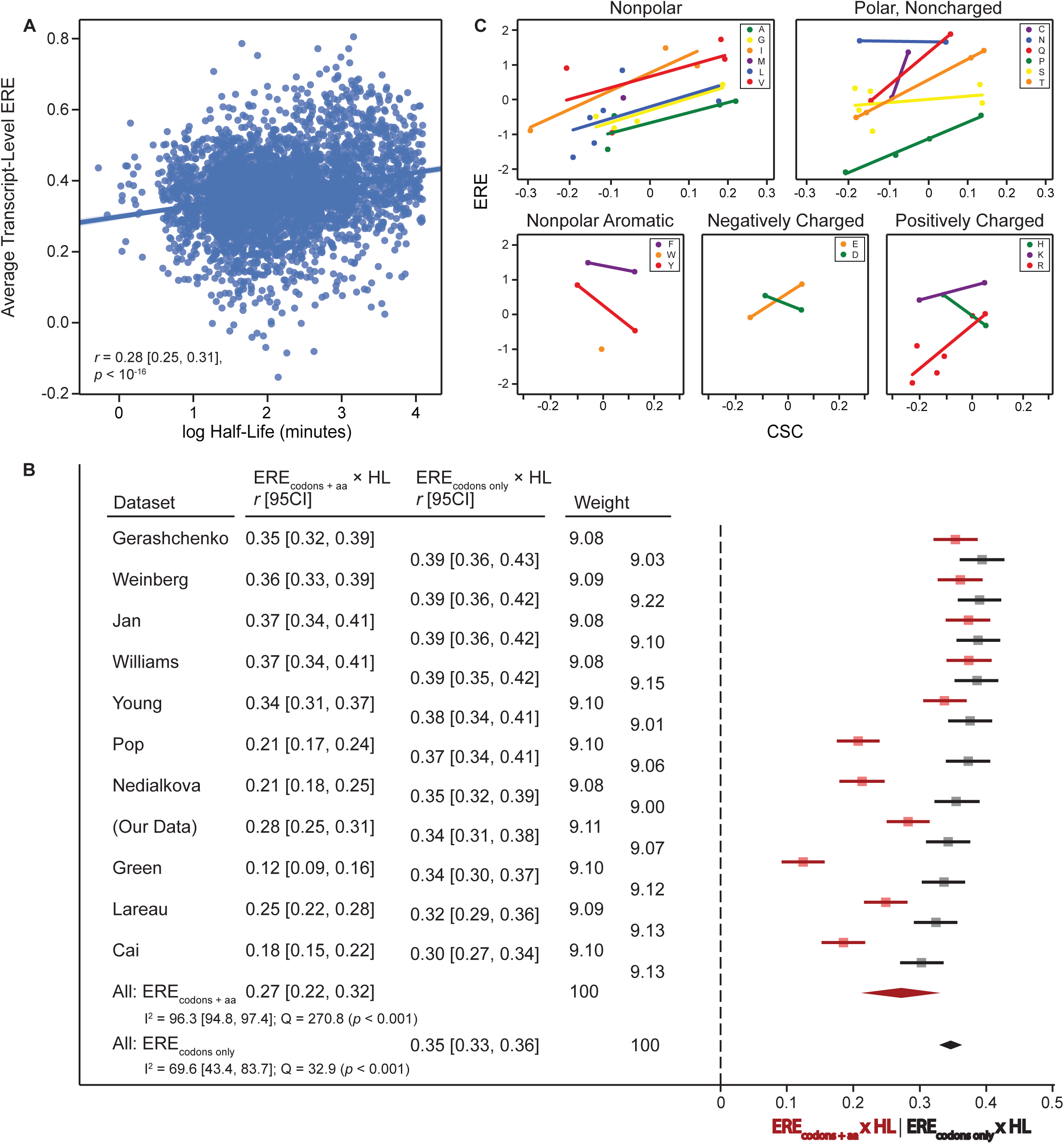
Ribosomal A-site decoding drives the relationship between elongation rate estimates and mRNA stability. *(A)* Scatter plot and best-fit line describing the relationship between average normalized elongation rate estimates (EREs) across a transcript and transcript stability. Transcript stability is plotted as the log of the transcript half-life, using data from (Presnyak et al. 2015). The correlation between transcript-level ERE and transcript half-life takes into account the per-gene variability in these estimates. *(B)* Forest plot describing the meta-analysis across 11 cycloheximide-minus ribosome profiling datasets of the correlation between the log of mRNA half-life and either transcript-level ERE (ERE_codons + aa_; red squares) or transcript-level ERE that takes into account the influence of codon identity, but not the influence of amino acid identity, on elongation rate (ERE_codons only_; black squares) with the lines corresponding to the 95% confidence interval. The aggregate correlation estimates for the ERE_codons + aa_ and ERE_codons only_ analyses are represented by the large red and black diamonds, respectively, with the width of the diamond corresponding to the 95% confidence interval of the aggregate estimate. I^2^ and Cochran’s Q are also reported as standard measures of heterogeneity. *(C)* Scatter plots of the relationship between normalized per-codon ERE and CSC, with grouping based on the properties of the encoded amino acid. The best-fit lines in each plot describe the relationship between ERE and CSC for each amino acid associated with more than 1 codon.

It has long been established that tRNA decoding is not the only step that can influence ribosome elongation dynamics. Different amino acids vary in the speed at which they are incorporated into a nascent polypeptide (Johansson et al. 2011), and certain amino acids are thought to have a significant impact on elongation (Wohlgemuth et al. 2008; Tanner et al. 2009; Watts and Forster 2010; Lareau et al. 2014). However, the observed relationship between tRNA abundance and CSC suggests that it is decoding that is driving the relationship between codon identity and mRNA stability rather than other aspects that affect elongation rate (i.e. amino acid identity). To distinguish these possibilities, we plotted the relationship between each codon’s ERE and CSC values after grouping the encoded amino acids based on their properties (Fig. 3C). From this analysis, we observe that individual amino acids span a wide range of elongation rates. For example, codons encoding proline and arginine exhibit some of the slowest elongation rates, as one might expect. However, it can readily be observed that for these two amino acids, synonymous codons still span a range of elongation rates that positively correlate with CSC (Fig. 3C). Considering all slopes of the correlation between ERE and CSC for all 18 amino acids decoded by more than one codon, the predominant trend is for these slopes to be positive (*t*(17) = 2.16, *p* < 0.05), showing that, in general, the faster the elongation rate for a given synonymous codon that encodes a certain amino acid, the higher the associated CSC value. Notably, five amino acids exhibit either no relationship between ERE and CSC for synonymous codons (asparagine) or a negative relationship (phenylalanine, tyrosine, aspartic acid, and histidine). While both tRNA decoding and amino acid identity are thought to influence translation elongation rate, these data suggest that tRNA decoding is primarily driving mRNA decay.

To explore this idea, we reasoned that if both decoding and amino acid contributions to elongation were influencing mRNA decay, then isolating decoding effects on translation elongation should decrease the correlation between transcript-level ERE and mRNA stability. To this end, we corrected the ERE metric calculated for each transcript such that it falls between 0 and 1, with 0 indicating the use of the slowest elongating codons at every amino acid position, and 1 indicating the use of the fastest elongating codons at every position (see Methods). This correction for amino acid identity results in the calculation of elongation rate estimates that only takes into account synonymous codon effects on elongation (ERE_codons only_), in contrast to our original elongation rate estimates that took into account both codon and amino acid effects on elongation (EREcodons + aa). Compared to the correlation we previously observed in our data between transcript-level ERE_codons + aa_ and mRNA half-life, the correlation between ERE_codons only_ and mRNA halflife is significantly higher (Fig. 3B; *r*= 0.34 compared to *r*= 0.28, *Δr*= 0.06, *N*=3969, *p*< 0.01), indicating that it is ribosomal A-site decoding that links translation elongation rates to mRNA decay rates.

The true impact of the amino acid correction is evident when we observe the effect of applying this correction to all *S*. *cerevisiae* datasets considered. While the correlation between ERE_codons + aa_ and transcript half-life in all datasets analyzed is 0.27, the correlation becomes even stronger when using ERE_codons only_ for our analysis (Fig. 3B; *black*). Interestingly, the relationship between ERE_codons only_ and transcript half-life shows significantly reduced between-dataset variability compared to the correlations observed with ERE_codons + aa_. This suggests that the primary source of variation between ribosome profiling datasets is in amino acid effects, and not in codon effects within a given amino acid. The clear consensus we observe between the datasets in support of a strong relationship between the amino acid-corrected ERE_codons only_ and mRNA stability is highly consistent with tRNA decoding rate being the primary component of elongation that determines mRNA stability in yeast.

### Ribosomal A-site decoding contributes to translation efficiency

Our global ribosome profiling analysis indicates that decoding of the codon in the ribosomal A-site, rather than encoded amino acid identity, impacts translation elongation rate and, subsequently, mRNA stability. While these genomic relationships are intriguing, we sought to validate these findings using a controlled system in which we use synonymous codons to manipulate codon optimality while maintaining the encoded amino acid sequence. To this end, we used a set of 11 *HIS3* reporters that vary from each other only in their percent codon optimality, from 0-100% in 10% increments (Fig. 4A). Importantly, by controlling for the encoded His3 protein sequence as well as 5’ and 3’ untranslated region sequences in each of these constructs, we can specifically study the impact of codon decoding on translation efficiency without interference from the effect that these parameters can have on translation rate. We previously used these *HIS3* reporters to demonstrate the positive correlation between codon optimality and mRNA stability, and we found that even small increases in codon optimality (e.g., 10%) can cause clearly detectable increases in mRNA stability (Radhakrishnan et al. 2016).

**Figure 4.**
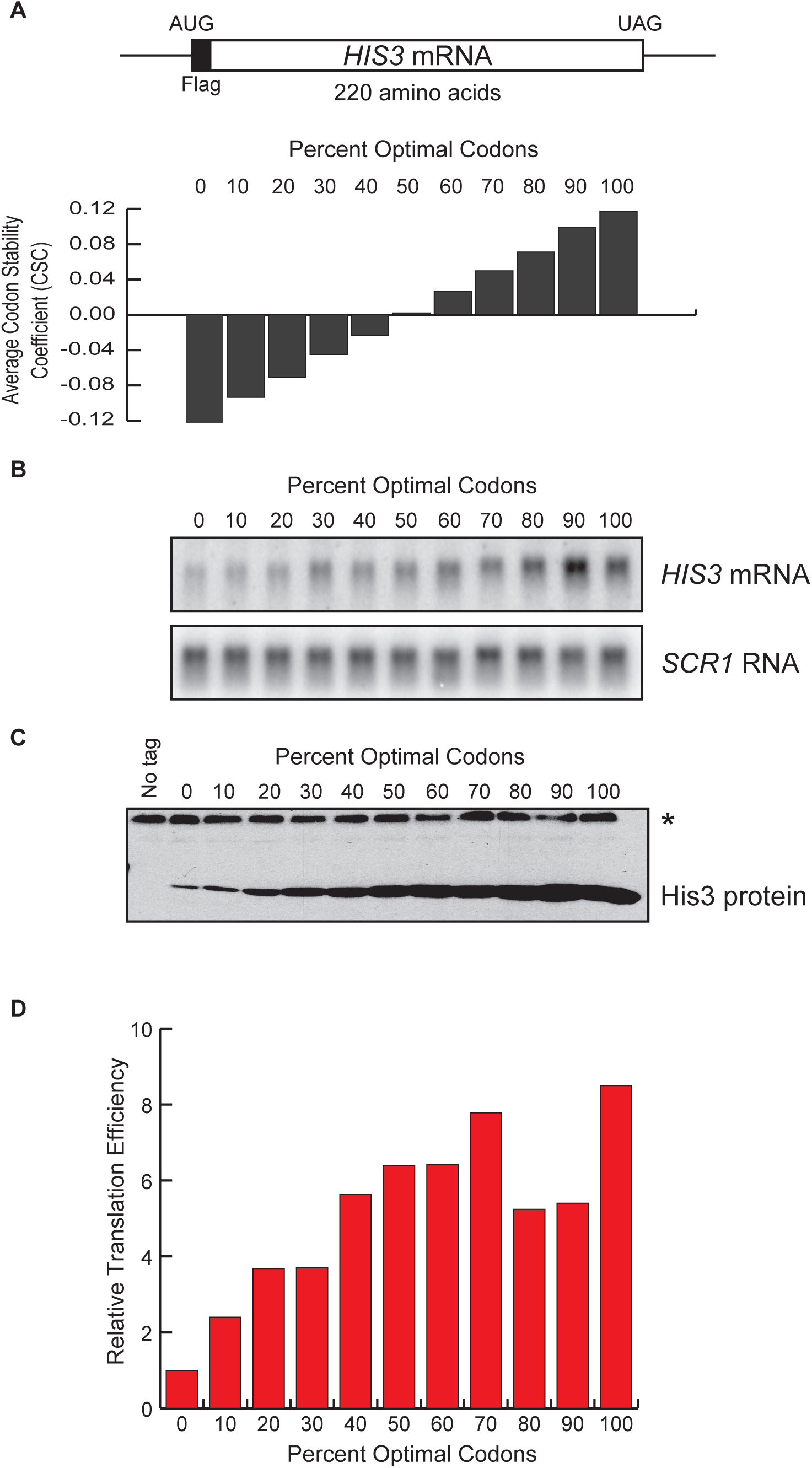
Translation efficiency is influenced by codon optimality. *(A)* Schematic representation of a *HIS3* mRNA with an N-terminal FLAG tag. The *HIS3* coding sequence was randomly altered using synonymous codons to generate constructs with varying percent codon optimality, from 0% to 100%. The average codon stability coefficient (CSC) for each construct is shown. CSC is a measure of the contribution of each codon to mRNA stability, and positive CSC values represent optimal codons while negative CSC values represent non-optimal codons. *(B)* Steady state *HIS3* mRNA levels expressed from the 0-100% optimal *HIS3* constructs were analyzed by Northern blot and were quantified relative to an *SCR1* loading control. *(C)* Steady state protein levels expressed from the 0-100% optimal *HIS3* constructs were analyzed by Western blot using anti-FLAG antibody. While the image presented here illustrates the clear correlation between codon optimality and steady state protein abundance using equivalent loading for each sample, more accurate quantification of protein levels was performed using variably diluted samples to minimize protein saturation of the higher optimality constructs. These more accurate values were used for the analysis shown in (Fig. 4D). The asterisk in this figure indicates the position of a non-specific band that is indicative of equal loading in all lanes. *(D)* Graphical representation of the translation efficiency of each *HIS3* construct relative to the 0% optimal construct. Translation efficiencies were determined by dividing the steady state protein abundance by the steady state mRNA abundance for each construct.

To test the effect of codon optimality on translation efficiency while eliminating amino acid and untranslated region-dependent effects, we performed Northern blot and Western blot analyses to measure the steady state *HIS3* mRNA and His3 protein levels expressed from each 0-100% optimality construct. We observed that whereas the steady state mRNA abundance varied up to ∼9-fold between each of the constructs (Fig. 4B), the steady state protein abundance differences were much more substantial (up to ∼49-fold; Fig. 4C). Due to the large range in steady state protein abundances that we observed for these constructs, and the relatively small dynamic range of Westerns, we present Fig. 4C for illustration purposes but performed protein abundance measurements after diluting samples to variable extents to minimize saturation effects. We calculated the protein output per mRNA as a measure of the translation efficiency of each construct and found that as percent codon optimality increases, translation efficiency generally increases (Fig. 4D). Interestingly, similar to our previous observation that even small differences in codon optimality cause detectable differences in mRNA stability, we see that even 10% differences in codon optimality also result in detectable differences in protein output per mRNA. The positive correlation between codon optimality and both translation efficiency and mRNA stability for this controlled set of constructs suggests that A-site decoding rates impact translation efficiency, and these differences in translation efficiency in turn affect the susceptibility of the mRNA to the decay machinery.

### Ribosomal A-site decoding contributes to translation kinetics

As an independent assay to test the influence of codon optimality on translation kinetics, we grew yeast cells expressing each of the 0-100% optimality *HIS3* constructs to mid-log phase before adding ^35^S-methionine/cysteine and harvesting the cells at different time points. The levels of ^35^S-methionine/cysteine-labeled His3 protein expressed from each construct at each time point were determined after running immunoprecipitated His3 protein on SDS-PAGE gels and detecting the ^35^S-labeled protein on a phosphorimager screen. To control for the effects of construct-dependent mRNA abundance or stability differences on the level of ^35^S-methionine/cysteine-labeled His3 protein produced, we internally normalized the data from each construct by calculating the amount of ^35^S-labeled His3 protein at each time point relative to the amount detected at the 2 minute time point for that construct.

In comparing the two optimality extremes (0% and 100%), we observed that the protein expression from the 0% optimality construct was substantially lower than the protein expression from the 100% optimality construct, consistent with what we observed in Fig. 4C (Fig. 5A). Importantly, the increase in protein expression from one time point to the next was much slower for the 0% optimality construct relative to the 100% optimality construct (Fig. 5A, B). Specifically, the rate of ^35^S incorporation was 2.8-fold slower for the 0% optimal construct than for the 100% optimal construct, indicating that translation of the higher optimality construct is more efficient (Fig. 5C). When analyzing the level of increase in the abundance of ^35^S-methionine/cysteine-labeled His3 protein over the time course for the remaining 9 constructs, we observed a general trend that was consistent with what was observed for the extremes, with relative ^35^S incorporation rates correlating with codon optimality (Fig. 5B, C). These data are in agreement with our previous observations and suggest that A-site decoding is linking translation kinetics to mRNA stability.

**Figure 5.**
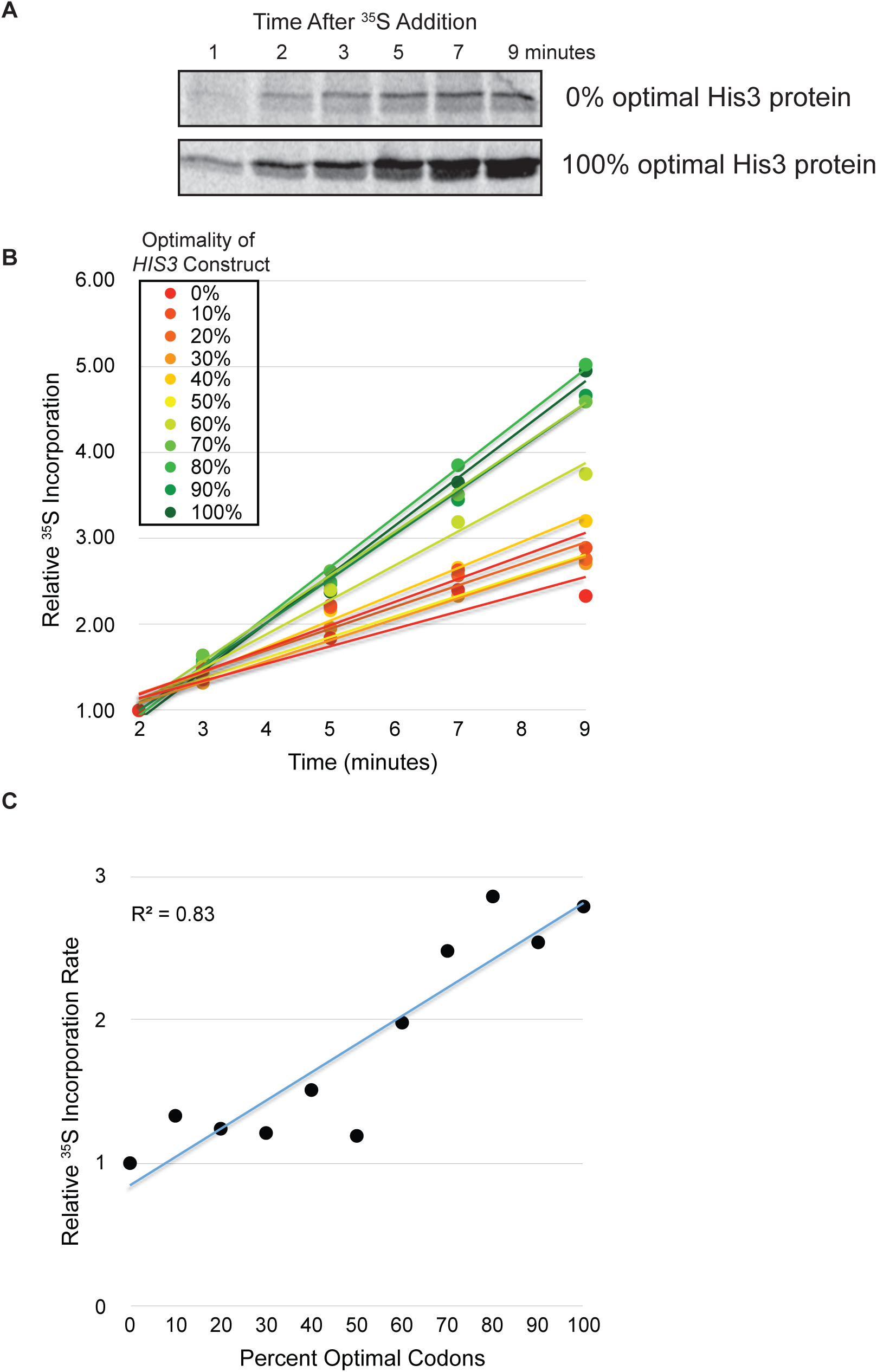
Codon optimality contributes to the rate of His3 protein production following ^35^S-methionine/cysteine labeling. *(A)* A representative image showing levels of ^35^S-methionine/cysteine-labeled His3 protein at each time point following addition of ^35^S-methionine/cysteine to *S*. *cerevisiae* cells at mid-log phase. His3 protein expressed from the 0% and 100% optimal *HIS3* constructs is presented. *(B)* Graphical representation of the increase in ^35^S-methionine/cysteine-labeled His3 protein abundance over time, relative to the protein abundance at the 2 minute time point, for each of the 0-100% optimal *HIS3* constructs. *(C)* The slope of each curve in Fig. 5B, normalized to the slope for the 0% optimal construct, is plotted as a measure of the relative rate of ^35^S incorporation per minute for each *HIS3* construct.

## Discussion

Previously, we had established that codon optimality is a powerful determinant of mRNA decay rates in the yeast *S. cerevisiae* (Presnyak et al. 2015). Other labs subsequently established this as a general principle across a number of species (Bazzini et al. 2016; Boël et al. 2016; Harigaya and Parker 2016; Mishima and Tomari 2016), leading to widespread interest in the general rules that dictate how translation elongation rates are communicated to the decay machinery. Our previous studies using reporter constructs suggest that decoding of codons in the A-site of the ribosome dictate elongation rate, which in turn sets mRNA decay rate via recruitment of the decapping factor Dhh1 (Radhakrishnan et al. 2016). However, whether codon optimality-mediated differences in translation elongation link directly to differences in mRNA stability has not been shown on a genome-wide scale.

We first performed ribosome profiling without cycloheximide in yeast as in (Hussmann et al. 2015; Weinberg et al. 2016) and were able to extract estimates of codon-specific elongation rates that correlate well with the tRNA adaptation index (tAI). Of note, these findings are in agreement with a number of ribosome profiling studies lacking cycloheximide (Hussmann et al. 2015; Weinberg et al. 2016). Further, codon-specific elongation rate estimates (EREs) correlate with codon occurrence to mRNA stability correlations (CSCs) that we calculated previously (Presnyak et al. 2015), suggesting that elongation rate at A-site codons genome-wide is influencing decay (Fig. 1C). Using our calculated per-codon ERE, we sought to determine how transcript-level elongation rates might influence decay. As would be predicted from codon-level elongation rates, transcripts with increasing proportions of optimal codons exhibit higher average EREs (Fig. 1D). Extending these data revealed that transcript-level EREs positively correlate with mRNA half-lives. Further analysis revealed that 10 previously published ribosome profiling datasets also yield a significant ERE by mRNA half-life correlation (Fig. 3B).

The correlations we observed between ERE and mRNA half-life in the 11 ribosome profiling datasets used in our study are statistically significant, but represent an aggregated correlation coefficient of only 0.27. This is not unexpected, as there are likely complex interactions between codons within open reading frames that are not accounted for by our modeling (Chevance et al. 2014; Gamble et al. 2016; Chevance and Hughes 2017). In addition, mRNA decay is not solely influenced by translation elongation, with clear contributions from other steps in translation and other mRNA features such as open reading frame length and cis-acting sequences in untranslated regions (Muhlrad and Parker 1992; Lee and Lykke-Andersen 2013; Geisberg et al. 2014; Neymotin et al. 2016). Together, these data extend previously observed correlations between codon-specific tAI and elongation rate by demonstrating that a statistically significant relationship exists between transcript-level elongation rate estimates and mRNA stability.

Translation elongation is a combination of decoding at the A-site as well as peptide bond formation rates that differ amongst amino acids (Watts and Forster 2010; Lareau et al. 2014). To uncover mechanistic features of how elongation might influence decay, we showed that EREs of synonymous codons generally positively correlate with the impact of codons on mRNA stability, independent of the encoded amino acid (Fig. 3C). Further, when we isolate synonymous codon effects on elongation from amino acid-specific effects, our correlations with mRNA half-life are improved (Fig. 3B). If amino acid-specific effects on elongation were contributing to mRNA decay rates, we would have seen a weakened correlation between codon-specific elongation rate (ERE_codons only_) and CSC (Fig. 3C). In addition, if amino acid identity substantially impacted mRNA decay rates, we would not have expected that most amino acids are encoded by synonymous codons that both contribute to the stabilization (positive CSC) and destabilization (negative CSC) of mRNAs. This highlights that A-site tRNA decoding is likely driving mRNA decay in yeast.

To rigorously test whether A-site decoding is affecting translation elongation which in turn affects decay, we utilized a set of 11 *HIS3* constructs that differ in total codon optimality from 0-100% in 10% increments. These constructs have the same untranslated regions, the same initiation context, and encode the exact same protein, making them an ideal system to isolate the effects of A-site decoding and to test incremental increase in the proportion of optimal codons. We had previously shown that the mRNAs expressed from these constructs exhibit an ∼13-fold range in half-lives, with an ∼3 minute half-life for the 0% optimal construct and an ∼40 minute half-life for the 100% optimal construct (Radhakrishnan et al. 2016). In the present study, we find a roughly 8.5-fold range of translation efficiency (protein per mRNA; Fig. 4) and a 2.8-fold range of relative ^35^S-methionine/cysteine incorporation rate across the *HIS3* constructs (Fig. 5), indicating differences in translation kinetics that are driven by decoding of codons. These data are in agreement with our previous observation that upon inhibiting translation initiation through glucose deprivation, existing ribosomes on the 100% optimal *HIS3* construct were cleared from the mRNA faster than ribosomes associated with the 0% optimal *HIS3* construct, presumably through more rapid elongation (Presnyak et al. 2015). While the codon-mediated effects on kinetics that we observed for this set of *HIS3* constructs are weaker than the range of translation efficiencies or half-lives observed, all measures trend in the same direction indicating that codon optimality drives changes in translation kinetics that in turn drive changes in mRNA decay rates.

This work makes two significant advances toward understanding the intimate relationship between mRNA translation and decay. First, we show that codon and transcript-level elongation rate estimates correlate with mRNA stability across the transcriptome. Second, we identify ribosomal A-site decoding as the step of elongation that impacts normal mRNA decay in yeast. In contrast, other work has suggested that both codon optimality and amino acid identity are important for influencing mRNA stability in zebrafish and *Xenopus* (Bazzini et al. 2016). These differences between yeast and higher eukaryotes may reflect differences in either the translation machinery, the decay machinery, or both. Alternatively, amino acid effects could be explained by the influence of amino acids on mRNA decay independent of translation elongation. For example, one possibility is that particular nascent amino acid sequences could recruit trans-acting factors that in turn regulate mRNA stability.

Strikingly, these findings are consistent with other known decay pathways: no-go, nonstop, and nonsense-mediated decay all are triggered by different states of the ribosomal A-site. In future work, it will be fascinating to determine how A-site decoding rates are transmitted to the normal mRNA decay machinery in yeast and also to determine how translation elongation influences decay in higher eukaryotes.

## Methods

### Yeast strains

The genotypes of the *S*. *cerevisiae* strains used in this study are listed in Supplemental Table S2. Yeast cells were grown to mid-log phase at 24°C in synthetic media (pH 6.5) containing 2% glucose and appropriate amino acids.

### Ribosome profiling library preparation

Using *S*. *cerevisiae* strain yJC2229, ribosome footprint RNA and control total RNA were isolated and libraries were prepared as was described in (Smith et al. 2014), with modifications. Specifically, cycloheximide treatment was omitted prior to cell harvest but was included during cell lysis. RNA purified from monosome fractions and control total RNA were depleted of ribosomal RNA once using the Yeast Ribo-Zero Gold rRNA Removal Kit (Illumina MRZY1324) according to the manufacturer’s instructions following the addition of an RNA Spike-In mix (Thermo Fisher Scientific 4456740) to the total RNA sample. cDNA libraries were amplified using the indexed primer oKB690 (Smith et al. 2014) and were sequenced at the Case Western Reserve University Genomics Core Facility using the Illumina HiSeq2500 platform.

### Ribosome profiling and RNA sequencing data processing

Adaptors were trimmed from the RNA and ribosome profiling datasets using cutadapt with the following parameters: -a CTGTAGGCACCATCAAT –trim-n –m 24 –M 36 –O 6. Other datasets were subject to adaptor trimming as appropriate using study-specific adaptor sequences, but otherwise identical trimming parameters. The ribosome profiling reads were then aligned against an index of *S*. *cerevisiae* ribosomal RNA sequences from Ensembl using bowtie with the following parameters: -D 15 –R 2-N 1 –L 25 –I S,1,0.75. Sequences that failed to align to the ribosomal RNA index were taken to be from messenger RNA and were aligned to the entire *S*. *cerevisiae* genome using HISAT2, with release 84 of Ensembl’s gene annotations of the sacCer3 genome (ftp://ftp.ensembl.org/pub/release-84/gtf/saccharomyces_cerevisiae/Saccharomyces_cerevisiae.R64-1-1.84.gtf.gz) to guide alignment to the transcriptome. Finally, multi-mapped reads were discarded and uniquely mapped ribosome footprint reads were transformed to transcriptome-based coordinates for further analysis using the sam2transcriptome python script. Ribosome footprint reads were then assigned to the A-site codon using the method outlined previously (Ingolia et al. 2009), where the P-site is identified based on the well characterized pileup of ribosome protected fragments over the start codon, with the P-site generally located 12 nucleotides into the fragments located at this start site peak, and the A-site another 3 nucleotides past this point. RNA-sequencing data was quantified with htseq-count (Anders et al. 2015) to estimate per-gene expression as reads per million mapped reads (RPM). For any ribosome profiling datasets without RNA-seq data, we used per-gene FPKMs calculated as the mean of all other comparable datasets’ RNA-seq.

### Model

Our primary aim was to extract estimates of the relative rates at which ribosomes move off of specific codons. Per-codon elongation rates have been estimated in the past (Qian et al. 2012; Pop et al. 2014; Hussmann et al. 2015; Weinberg et al. 2016). However, the common approach to estimating codon-specific elongation times has been to simply calculate the ribosome density over a codon relative to the frequency of a codon in an mRNA, and then sum these values up across the entire transcriptome. This approach is intuitive and easily calculated, but it is difficult to assess the confidence of the estimates generated from this procedure. There is often no control for biases introduced by those genes with sparse ribosome protected fragment coverage, which are weighted equally with the rest of the transcriptome in the default approach.

To extract more robust estimates of codon-specific ribosome elongation rates, we propose the following model, which is based on the assumption that ribosome initiation is typically a much slower process than elongation (Shah et al. 2013). Let *ρ_g_* represent the initiation rate for a gene *g*, and *τ_c_* be the average elongation time for a codon type *c* ∈ {*AAA*,*AAC*,*AAG*,…,*TTT*}. Thus, under our assumption, for a single mRNA molecule from gene *g*, the proportion of time that a ribosome can be found on a position with codon *C* is:

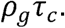

Let the relative concentration, as measured by RNA-seq reads mapped to gene *g* per million mapped reads for gene *g* be denoted by *M_g_*, and let the number of *c* codons in gene *g* be *N*_*g*,*c*_. Also, let *Y*_*g*,*c*_ be the number of ribosome footprints mapping to *c* codons in gene *g*. Based on this, we assume that *Y*_*g*,*c*_ follows an overdispersed Poisson distribution with the following mean:

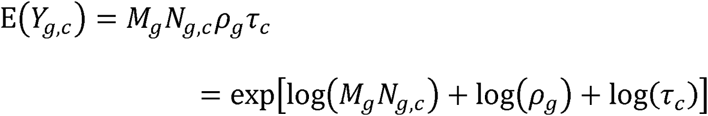

An overdispersed Poisson distribution was chosen to model ribosomal reads as it has previously been found that the sampling rate of RNA fragments from their associated transcripts can rarely be described as following a Poisson distribution due to uneven scaling of variability with the mean (Soneson and Delorenzi 2013). It must be understood that there is a degeneracy in this model; there are an infinite number of parameters which fit the data equally well. However, we can estimate the values *τ_c_* relative to *ρ_g_*. To estimate the model parameters, we used a generalized linear mixed model which assumes that log(*ρ_g_*) ∼*Normal*(0,*s*^2^) where *s*^2^ and log(*τ_c_*) are all estimated from the data using maximum likelihood with an overdispersed Poisson distribution. The mixed model gets around the degeneracy (i.e. identifiability problem) by assuming that on average across all genes E[log(*ρ_g_*)] = 0. However, it must be understood that the resultant parameter estimates (*ρ̂_g_* and *τ̂_c_*) are relative. Given that estimates of codon-specific elongation times are relative, we will adopt the convention of first calculating the relative elongation rate
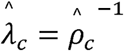 and then normalizing the resultant rate estimates so that E
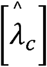
= 0 and *s*^2^= 1 for ease of interpretability. This model implicitly more highly weights those transcripts with a greater density of information on ribosome dynamics, removing potential biases due to sparse ribosome protected fragment density and avoiding the need to set an arbitrary threshold for ribosome coverage in an attempt to mitigate these effects. However, because the model can be easily represented as a linear mixed effects model, it can be estimated directly from ribosome profiling data using out of the box solvers in MATLAB or R, making it readily applicable to a variety of datasets.

### Ribosome profiling and RNA sequencing data analysis

For each gene, we first find the number of codons *c* (*N*_*g*,*c*_) in gene *g*, and also calculate the total number of ribosome protected fragments over gene *g* with the A-site mapped to codon *c* (*Y*_*g*,*c*_). Relative RNA concentration (*M_g_*) is calculated using htseq-count and standardized as RPM, as outlined above. This procedure is performed for each gene in the dataset, with no restriction on total ribosome density or gene length, though only genes with non-zero RNA expression and RPF counts are included. Data from between 4188 and 5245 genes were included, depending on the dataset analyzed, ensuring that we are able to leverage the majority of protein coding genes for analysis. For the model, data are structured as long-form arrays, with the number of rows equal to the <u>product</u> of the number of genes considered <u>times</u> the number of amino acid-coding codons (61), and the number of columns equal to 5 (codon identity, gene identity, number of ribosome protected fragments over a given codon in a gene, the total occurrence of that codon in a gene, and the estimate of gene expression). The model itself is fit using the *fitglme* function in the Statistical Toolbox of MATLAB, version R2016a. The code used to form ribosome profiling data into appropriate datasets, and the MATLAB code used to specify and run the model is available as supplementary information. All other statistical analyses were carried out using the statsmodels and scipy packages for Python, and Matplotlib and Seaborn were used to generate Fig. 1-3. The method used to calculate the statistical significance of a change in Pearson correlation coefficient was taken from (Cohen et al. 2013). Briefly, the test statistic for the change in the Pearson correlation coefficient can be found as follows:

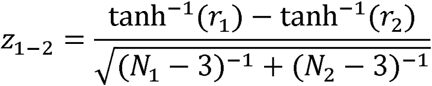

where *N_i_* is the number of observations used to calculate *r_i_*. This statistic is drawn from a normal distribution, and the corresponding p-value can be found with the survival function of a normal distribution with *μ* = 0 and *σ* = 1.

### Calculation of robust Pearson correlation coefficients

The standard method of calculating Pearson correlation coefficients has no facility to take into account the precision associated with the individual quantities that make up vectors **X** and **Y**, the two vectors for which the correlation is to be calculated. Therefore, we chose to leverage the modeling capabilities of Stan (Carpenter et al. 2017) to specify a model where **X** and **Y** are latent variables sampled from a Gaussian distribution such that **X** ∼ *Normal*(**X, Λ_1_**) and **Y** ∼ *Normal*(**Y,Λ_2_**), where **X** and **Y** are the point estimates for each quantity, and Λ_1_ and Λ_2_ are the standard errors associated with the values in **X** and **Y**, respectively. **X** and **Y** may be CSC values and ribosome elongation rate estimates, respectively, or gene-level average elongation rates and mRNA half-lives, depending on the needs of the analysis. Finally, we find the value of *r*, the Pearson correlation coefficient, to maximize the likelihood of the latent variables **X** and **Y** as samples drawn from a multivariate Gaussian distribution with means *μ_X_*,*μ_Y_* and the covariance matrix described as follows:

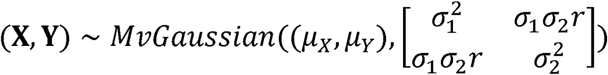

where *σ*_1_ and *σ*_2_ are the standard deviations of **X** and **Y**, respectively, and are also estimated in the model. The resultant sampled *r* values then represent the maximum a posteriori (MAP) distribution of *r*. The code to accomplish this was adapted from (Lee and Wagenmakers 2014; Schwarzer and Carpenter 2015). All models in Stan were run with 1000 warmup samples and 1000 acquisition samples, across four separate chains. Models were assessed for convergence by ensuring that the scale reduction factor across chains, *R̂*, was equal to 1.

### Meta-analysis

The meta-analysis of Pearson correlation coefficients follows the Inverse Variance method for calculating the random effects estimate *θ_R_*, using the DerSimonian-Laird estimator for calculating the between-study variability parameter *τ*^2^, and employing the Hartung and Knapp correction when estimating **var**(*θ_R_*). All meta-analyses were implemented in Python using custom code based on the methods presented in (Schwarzer and Carpenter 2015). Briefly, let the population estimate of the correlation between CSC and translation elongation rate (or average elongation rate and mRNA half-life), *r*, be expressed as the normally distributed Fisher’s *z* = tanh^−1^(*r*), with the point estimate and variance of *z* estimated from the a posteriori distribution of *r* calculated from Stan. With *z* sampled from a normal distribution with known variance, for each dataset we can define individual estimates of *z* for dataset *k* as:

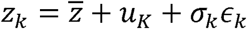

Where *Z* is the true transformed correlation coefficient in the population of datasets, *u_k_* ∼ *N*(0,*τ*^2^) captures the error due to heterogeneity between studies, while *ϵ_k_* ∼ *N*(0,1), scaled by the known standard error of
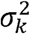
= **var**(*z_k_*), represents within-study error. The estimate of *Z* is simply the weighted average of individual estimates *Z_k_*, where the weight with which each study is considered in the analysis,
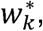
is inversely proportionate to the sum of the study-specific variance plus the between-study variance.
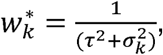leaving us with

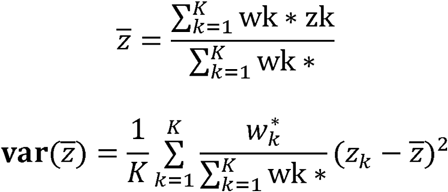

In order to calculate the 95% confidence interval around *z̅*, take *z̅* ±
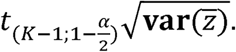
To express the results of this analysis is terms of *r̅*, simply take the hyperbolic tangent of *Z* and the upper and lower bounds of *z* calculated above.

### Calculation of amino acid-corrected elongation rate

An imaginary polypeptide can be perfectly efficient if every amino acid is coded with the fastest elongating codon, or perfectly inefficient if every amino acid is coded with the slowest elongating codon. Any further changes would require altering the amino acid sequence. We wished to calculate the elongation rate of a natural transcript relative to these two extremes, in order to remove effects of amino acid choice and effectively correct for any biases due to a transcript being enriched in amino acids that happen to be associated with faster or slower total ribosome transit times, regardless of the decoding speed. To accomplish this, we use elongation rate estimates to find the maximum and minimum elongation rate associated with each amino acid. Then, for a given transcript, we calculate the hypothetical maximum (*ER_max_*) and minimum (*ER_min_*) average elongation rates given the amino acid sequence associated with the transcript. The correction is then calculated as follows: F(max^ER^) =
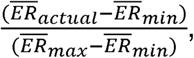
where *ER_actual_* is the average elongation rate based on the actual codon sequence of the transcript.

### Northern and Western blot analysis

*S*. *cerevisiae* cells expressing N-terminally FLAG-tagged 0-100% optimal *HIS3* (yJC2419 to yJC2429), or an untagged 100% optimal *HIS3* control (yJC2088), were used for total RNA or protein isolation as was described in (Geisler et al. 2012). Equal amounts of each RNA were run on 1.4% agarose formaldehyde gels, transferred onto nylon membrane, and probed using a ^32^P end-labeled oligonucleotide (oJC2564) (Presnyak et al. 2015), which is complementary to a 23 nucleotide region within each *HIS3* construct that was maintained for detection purposes. For protein analysis, either an equal amount of each sample, or variable amounts of each sample (in order to enable more accurate protein abundance quantification by minimizing signal saturation), were run on SDS-PAGE gels and were transferred onto PVDF membrane. His3 protein was detected using rabbit anti-FLAG primary antibody (Sigma F7425) and goat antirabbit secondary antibody (Pierce 31460). Steady state RNA and protein abundance was quantified using ImageQuant and ImageJ software, respectively. Translation efficiency was determined by dividing the steady state His3 protein abundance by the steady state *HIS3* mRNA abundance for each construct.

### ^35^S-methionine/cysteine incorporation and FLAG-His3 protein immunoprecipitation assay

*S*. *cerevisiae* cells expressing N-terminally FLAG-tagged 0-100% optimal *HIS3* (yJC2488 to yJC2498), were grown in the absence of supplemented methionine and cysteine. At mid-log phase, the cells were concentrated 10-fold by pelleting and resuspending in the same media used for growth. Next, 15 μL of ^35^S-methionine/cysteine mix (Perkin Elmer NEG072) per 50 mL of cells were added to each culture and cell aliquots were transferred into an equal volume of 2X Buffer A (50 mM sodium azide, 100 mM NaCl, 10 mM Tris-HCl pH 7.4, 5 mM MgCl_2_, 5 mM NH_4_Cl, 1 mM DTT, 2 μL/mL protease inhibitor (Sigma p8215), 200 μg/mL cycloheximide) on ice at the time points indicated in Fig. 5A. Cells were pelleted, washed, and then lysed in 1X Buffer A using glass beads and alternating cycles of vortexing and incubating on ice. Lysates were quantified and an equivalent amount of each time point within a time course (typically between 25 and 40 OD_260_ units) were incubated with rabbit anti-FLAG antibody (13.3 μL/mL) in 1X Buffer A supplemented with NP40 to 0.1% and 3 μL/mL protease inhibitor for 1 hour 10 min. at 4°C. Anti-FLAG antibody-bound His3 protein was then immunoprecipitated following a 1 hour incubation with Dynabeads Protein G (Invitrogen 10004D). After removal of the supernatant, the Dynabeads were washed 5 times with IP Wash Buffer (150 mM Tris-HCl pH 7.5, 125 mM NaCl, 2 mM MgCl_2_, 0.1% NP40) and FLAG-His3 protein was eluted by heating the beads at 95°C for 5 min. in IP Elution Buffer (50 mM Tris-HCl pH 7.5, 0.5% SDS, and 50 mM EDTA pH 8.0). Eluates were run on SDS-PAGE gels, transferred onto PVDF membrane, and then exposed to a phosphorimager screen. Detected ^35^S-labeled His3 protein was quantified using ImageQuant software and normalized to the 2 min. time point for each time course.

## Declarations

### Ethics approval and consent to participate

Not applicable

### Consent for publication

Not applicable

### Availability of data and materials

The datasets generated and/or analyzed during the current study are available in the Gene Expression Omnibus. Details and GEO numbers for each dataset are in Supplemental Table S1.

### Authors’ contributions

GH and NM developed the statistical models detailed in the Methods section for analyzing ribosome profiling data. NA performed experiments for figures 1-5. GH performed the analyses for figures 1-3. GH, NA, TS, and JC wrote the manuscript and all authors read and approved the final manuscript.

## Acknowledgements

We thank all members of the Coller lab and the lab of Dr. Kristian Baker for helpful comments and discussion. We also thank DaJuan Whiteside for technical assistance.

---

**Supplemental Figure S1.** Comparison of elongation rate estimates from 11 S. *cerevisiae* datasets. Heat map of elongation rate estimates from 11 datasets, standardized within each dataset to allow easy comparison of rates across datasets. This heat map reveals strong groupings of codons into relatively fast (red) and slow (blue) sets that hold across most datasets analyzed. On the right, the correlation matrix of elongation rates across all datasets is presented, showing the strong positive correlations between many datasets, as well as highlighting those datasets that are less well correlated with the others. Pearson correlation coefficients are encoded in the color of the cells in the matrix, as well as in the printed text within each cell.

## Supplemental Methods

### Ribosome profiling and RNA sequencing data processing

Adaptors were trimmed from the RNA and ribosome profiling datasets using cutadapt with the following parameters: -a CTGTAGGCACCATCAAT –trim-n –m 24 –M 36 –O 6. Other datasets were subject to adaptor trimming as appropriate using study-specific adaptor sequences, but otherwise identical trimming parameters. The ribosome profiling reads were then aligned against an index of S. ***cerevisiae*** ribosomal RNA sequences from Ensembl using bowtie with the following parameters: -D 15 –R 2-N 1 –L 25 –I S,1,0.75. Sequences that failed to align to the ribosomal RNA index were taken to be from messenger RNA and were aligned to the entire S. ***cerevisiae*** genome using HISAT2, with release 84 of Ensembl’s gene annotations of the sacCer3 genome (ftp://ftp.ensembl.org/pub/release-84/gtf/saccharomyces_cerevisiae/Saccharomyces_cerevisiae.R64-1-1.84.gtf.gz) to guide alignment to the transcriptome. Finally, multi-mapped reads were discarded and uniquely mapped ribosome footprint reads were transformed to transcriptome-based coordinates for further analysis using the sam2transcriptome python script. Ribosome footprint reads were then assigned to the A-site codon using the method outlined previously (Ingolia et al. 2009), where the P-site is identified based on the well characterized pileup of ribosome protected fragments over the start codon, with the P-site generally located 12 nucleotides into the fragments located at this start site peak, and the A-site another 3 nucleotides past this point. RNA-sequencing data was quantified with htseq-count (Anders et al. 2015) to estimate per-gene expression as reads per million mapped reads (RPM).

### Model to estimate elongation rate

To extract more robust estimates of codon-specific ribosome elongation rates, we propose the following model, which is based on the assumption that ribosome initiation is typically a much slower process than elongation (Shah et al. 2013). Let *ρ_g_* represent the initiation rate for a gene *g*, and *τ_C_* be the average elongation time for a codon type *C* ( {*AAA*, *AAC*, *AAG*,…, *TTT*}. Thus, under our assumption, for a single mRNA molecule from gene ***g*,** the proportion of time that a ribosome can be found on a position with codon *c* is:

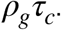

Let the relative concentration, as measured by RNA-seq reads mapped to gene *g* per million mapped reads for gene *g* be denoted by ***M**_g_*, and let the number of *c* codons in gene *g* be ***N***_*g*,*C*._ Also, let ***Y***_*g*,*C*_ be the number of ribosome footprints mapping to ***c*** codons in gene g. Based on this, we assume that ***Y***_*g*,*c*_ follows an overdispersed Poisson distribution with the following mean:

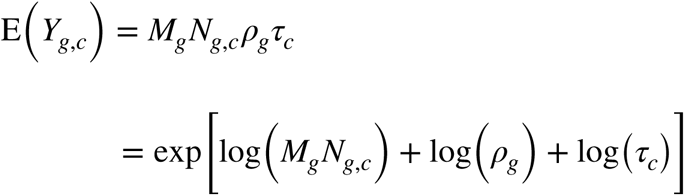

An overdispersed Poisson distribution was chosen to model ribosomal reads as it has previously been found that the sampling rate of RNA fragments from their associated transcripts can rarely be described as following a Poisson distribution due to uneven scaling of variability with the mean (Soneson and Delorenzi 2013). It must be understood that there is a degeneracy in this model; there are an infinite number of parameters which fit the data equally well.

However, we can estimate the values *τ_C_* relative to *ρ_g_*. To estimate the model parameters, we used a generalized linear mixed model which assumes that log
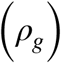
-*Normal* (0,*s*^2^) where *s*^2^ and log(*τ_C_*) are all estimated from the data using maximum likelihood with an overdispersed Poisson distribution. The mixed model gets around the degeneracy (i.e. identifiability problem) by assuming that on average across all genes
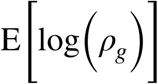
= 0. However, it must be understood that the resultant parameter estimates (*ρ̂_g_* and *τ̂_C_*) are relative. Given that estimates of codon-specific elongation times are relative, we will adopt the convention of first calculating the relative elongation rate
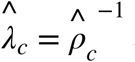
and then normalizing the resultant rate estimates so that
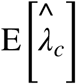
=0 and *s*^2^ = 1, for ease of interpretability.

This model implicitly more highly weights those transcripts with a greater density of information on ribosome dynamics, removing potential biases due to sparse ribosome protected fragment density and avoiding the need to set an arbitrary threshold for ribosome coverage in an attempt to mitigate these effects. However, because the model can be easily represented as a linear mixed effects model, it can be estimated directly from ribosome profiling data using out of the box solvers in MATLAB or R, making it readily applicable to a variety of datasets.

### Ribosome profiling and RNA sequencing data analysis

For each gene, we first find the number of codons *c* (***N***_*g*,*c*_) in gene *g*, and also calculate the total number of ribosome protected fragments over gene *g* with the A-site mapped to codon *c* (***Y***_*g*,*c*_). Relative RNA concentration (***M***_*g*,*c*_) is calculated using htseq-count and standardized as RPM, as outlined above. This procedure is performed for each gene in the dataset, with no restriction on total ribosome density or gene length, though only genes with non-zero RNA expression and RPF counts are included. Data from between 4188 and 5245 genes were included, depending on the dataset analyzed, ensuring that we are able to leverage the majority of protein coding genes for analysis. For the model, data are structured as long-form arrays, with the number of rows equal to the <u>product</u> of the number of genes considered <u>times</u> the number of amino acid-coding codons (61), and the number of columns equal to 5 (codon identity, gene identity, number of ribosome protected fragments over a given codon in a gene, the total occurrence of that codon in a gene, and the estimate of gene expression). The model itself is fit using the *fitglme* function in the Statistical Toolbox of MATLAB, version R2016a. The code used to form ribosome profiling data into appropriate datasets, and the MATLAB code used to specify and run the model is available as supplementary information. All other statistical analyses were carried out using the statsmodels and scipy packages for Python, and Matplotlib and Seaborn were used to generate Fig. 1-3. The method used to calculate the statistical significance of a change in Pearson correlation coefficient was taken from (Cohen et al. 2013). Briefly, the test statistic for the change in the Pearson correlation coefficient can be found as follows:

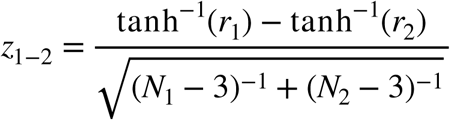

where ***N_i_*** is the number of observations used to calculate ***r_i_***. This statistic is drawn from a normal distribution, and the corresponding p-value can be found with the survival function of a normal distribution with ***μ*** =0 and ***σ*** = 1.

### Calculation of robust Pearson correlation coefficients

The standard method of calculating Pearson correlation coefficients has no facility to take into account the precision associated with the individual quantities that make up vectors **X** and **Y**, the two vectors for which the correlation is to be calculated. Therefore, we chose to leverage the modeling capabilities of Stan (Carpenter et al. 2017) to specify a model where **X** and **Y** are latent variables sampled from a Gaussian distribution such that **X** ∼ *Normal*(**X**, **Λ_1_**) and **Y** ∼ *Normal*(**Y**, **Λ_2_**), where **X** and **Y** are the point estimates for each quantity, and **Λ_1_** and **Λ_2_** are the standard errors associated with the values in **X** and **Y**, respectively. **X** and **Y** may be CSC values and ribosome elongation rate estimates, respectively, or gene-level average elongation rates and mRNA half-lives, depending on the needs of the analysis. Finally, we find the value of *r*, the Pearson correlation coefficient, to maximize the likelihood of the latent variables **X** and **Y** as samples drawn from a multivariate Gaussian distribution with means ***μ_X_*, *μ_Y_*** and the covariance matrix described as follows:

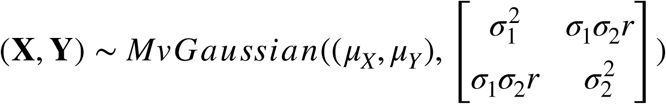

where *σ*_1_ and *σ*_2_ are the standard deviations of **X** and **Y**, respectively, and are also estimated in the model. The resultant sampled *r* values then represent the maximum a posteriori (MAP) distribution of r. The code to accomplish this was adapted from (Lee et al. 2014). All models in Stan were run with 1000 warmup samples and 1000 acquisition samples, across four separate chains. Models were assessed for convergence by ensuring that the scale reduction factor across chains, *R̂*, was equal to 1.

### Meta-analysis

The meta-analysis of Pearson correlation coefficients follows the Inverse Variance method for calculating the random effects estimate *θ_R_*, using the DerSimonian–Laird estimator for calculating the between-study variability parameter *τ*^2^, and employing the Hartung and Knapp correction when estimating **var**(*θ_R_*). All meta-analyses were implemented in Python using custom code based on the methods presented in (Schwarzer et al. 2015). Briefly, let the population estimate of the correlation between CSC and translation elongation rate (or average elongation rate and mRNA half-life), *r*, be expressed as the normally distributed Fisher’s *z* = tanh^−l^(*r*), with the point estimate and variance of *z* estimated from the a posteriori distribution of *r* calculated from Stan. With *z* sampled from a normal distribution with known variance, for each dataset we can define individual estimates of *z* for dataset *k* as:

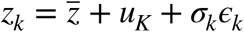

Where *z* is the true transformed correlation coefficient in the population of datasets, ***u_k_*** ∼ ***N***(0,T^2^) captures the error due to heterogeneity between studies, while ***ϵ_k_*** ∼ ***N***(0,1), scaled by the known standard error of
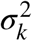
= **var**(*z_k_*), represents within-study error. The estimate of *z* is simply the weighted average of individual estimates *z_k_*, where the weight with which each study is considered in the analysis,
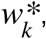
is inversely proportionate to the sum of the study-specific variance plus the between-study variance.
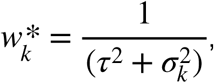leaving us with

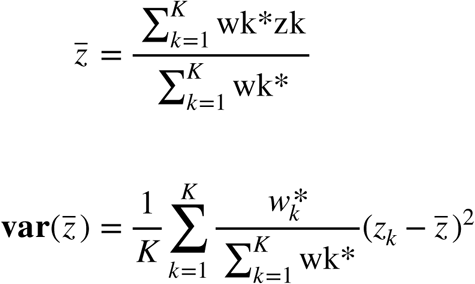

In order to calculate the 95% confidence interval around *z*, take *z* ±
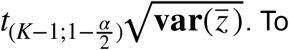
To express the results of this analysis is terms of *r*, simply take the hyperbolic tangent of *z* and the upper and lower bounds of *z* calculated above.

### Calculation of amino acid-corrected elongation rate

An imaginary polypeptide can be perfectly efficient if every amino acid is coded with the fastest elongating codon, or perfectly inefficient if every amino acid is coded with the slowest elongating codon. Any further changes would require altering the amino acid sequence. We wished to calculate the elongation rate of a natural transcript relative to these two extremes, in order to remove effects of amino acid choice and effectively correct for any biases due to a transcript being enriched in amino acids that happen to be associated with faster or slower total ribosome transit times, regardless of the decoding speed. To accomplish this, we use elongation rate estimates to find the maximum and minimum elongation rate associated with each amino acid. Then, for a given transcript, we calculate the hypothetical maximum (*ER_max_*) and minimum (*ER_min_*) average elongation rates given the amino acid sequence associated with the transcript. The correction is then calculated as follows:
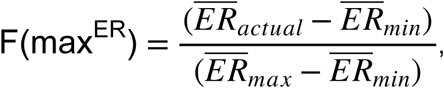 where *ER_actuai_* is the average elongation rate based on the actual codon sequence of the transcript.

## Supplementary Figure Legends

**Supplementary Figure S1:**
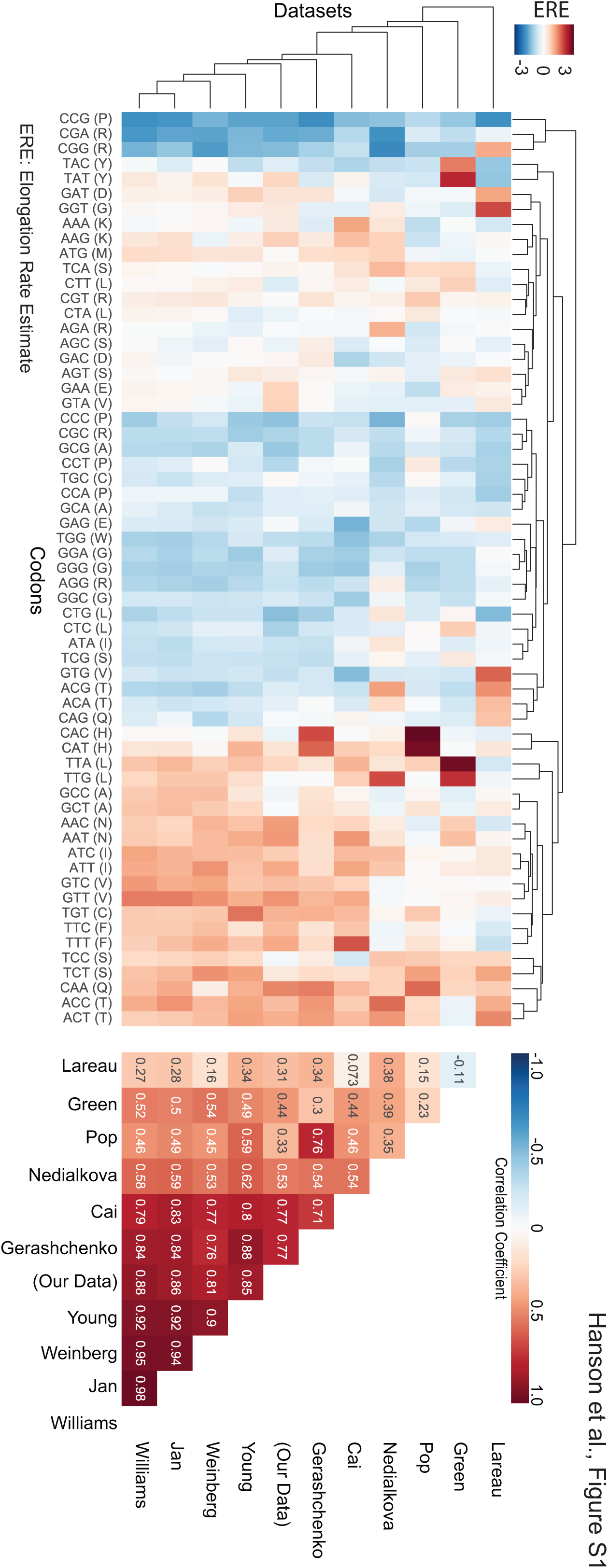
Heat map of elongation rate estimates from 11 datasets. Heat map of elongation rate estimates from 11 datasets, standardized within each dataset to allow easy comparison of rates across datasets. This heat map reveals strong groupings of codons into relatively fast (red) and slow (blue) sets that hold across most datasets analyzed. On the right, the correlation matrix of elongation rates across all datasets is presented, showing the strong positive correlations between many datasets, as well as highlighting those datasets that are less well correlated with the others. Pearson correlation coefficients are encoded in the color of the cells in the matrix, as well as in the printed text within each cell.

---

**Table S1:**
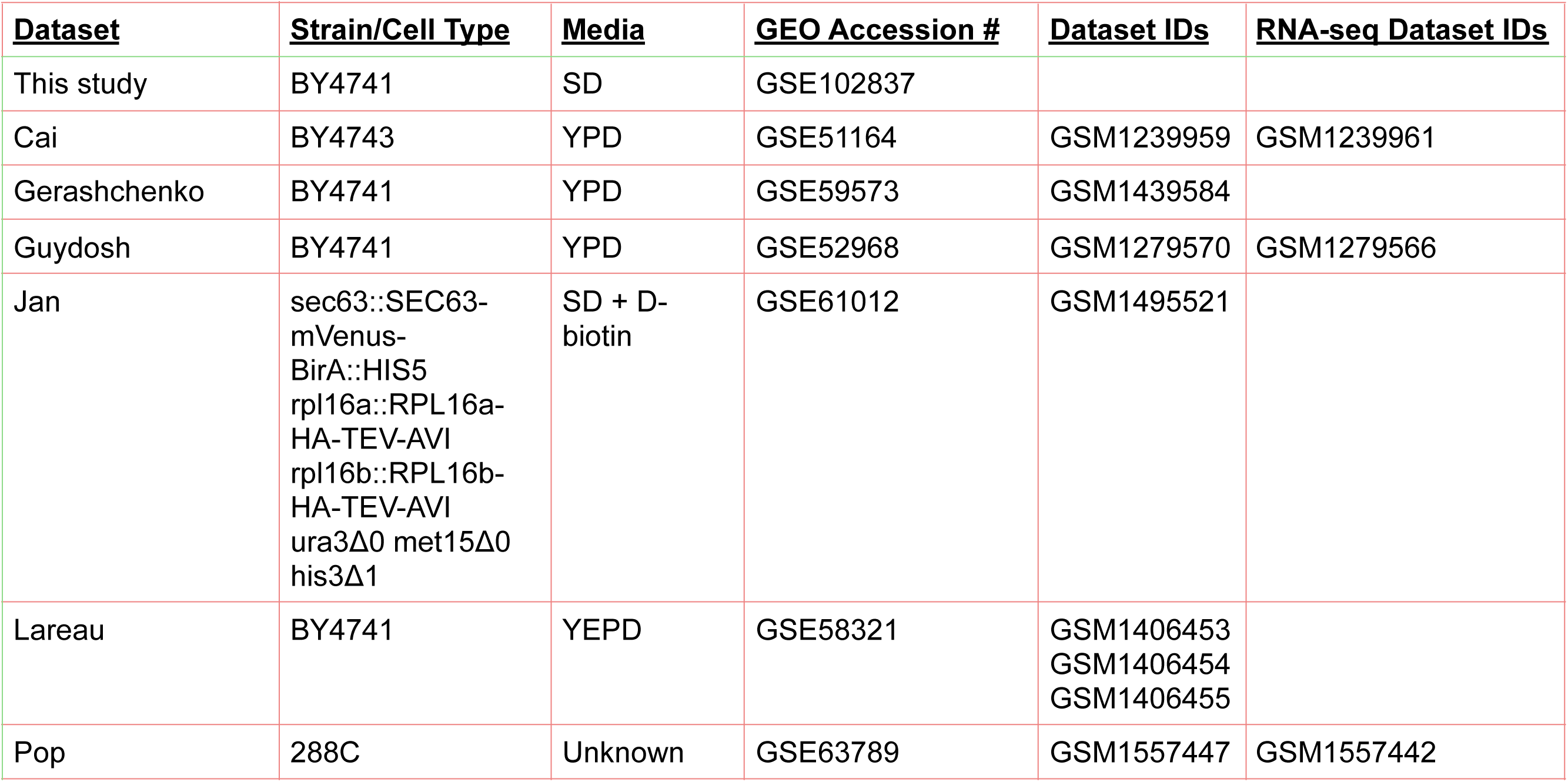
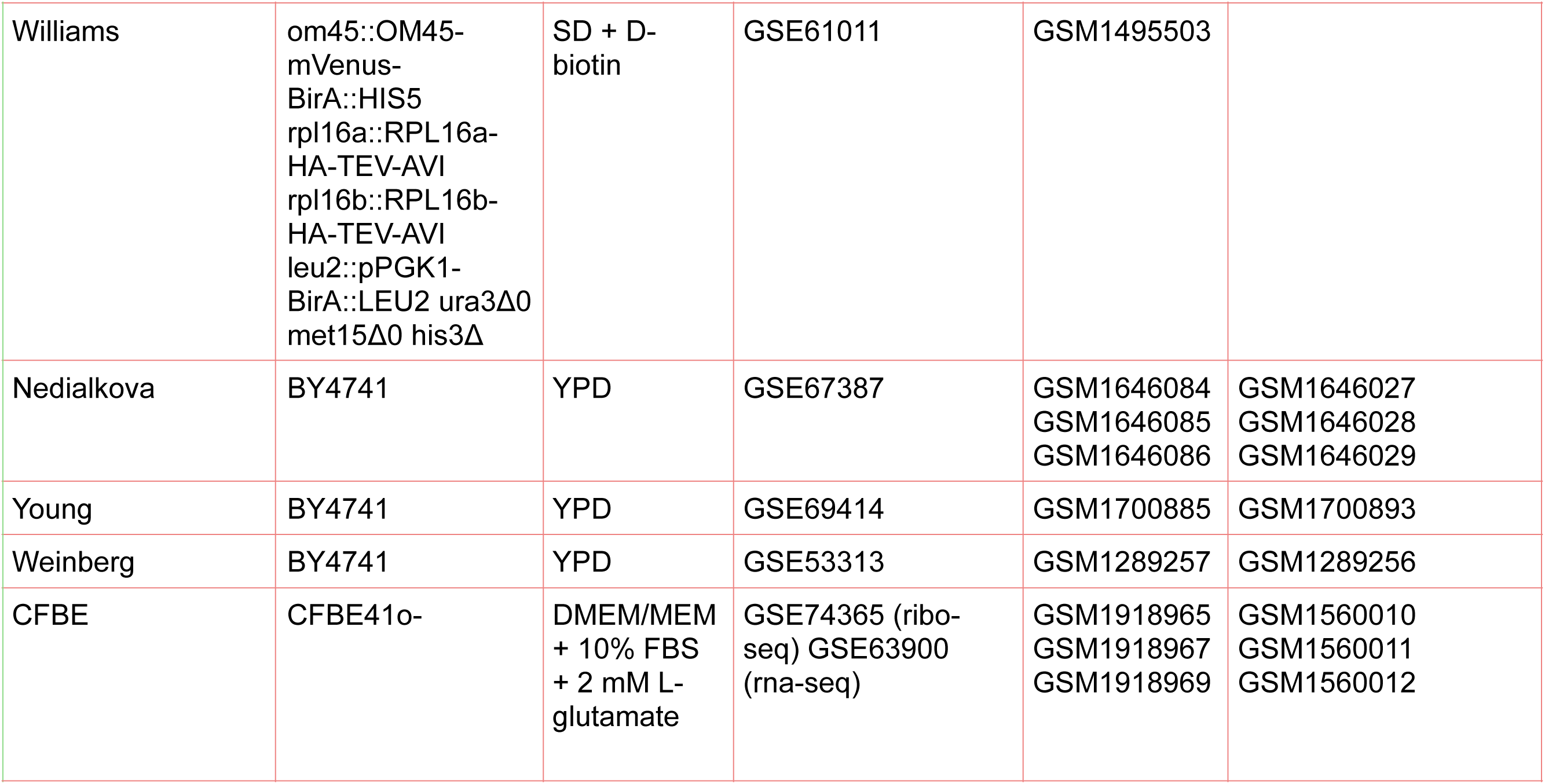
Datasets.

**Table S2:** Yeast strains.

| **<u>Strain name</u>** | **<u>Genotype</u>** | **<u>Source</u>** |
| --- | --- | --- |
| yJC2088 | MATa, *ura3*, *leu2*, *his3*, *met15*, [100% optimal *HIS3*, *URA3*] | This study |
| yJC2229 | MATa, *ura3*, *leu2*, *his3*, *met15*, [100% optimal *HIS3*, *URA3*], [0% optimal *HIS3*, *LEU2*] | This study |
| yJC2419 | MATa, *ura3*, *leu2*, *his3*, *met15*, [N-terminally FLAG tagged 0% optimal *HIS3*, *URA3*] | This study |
| yJC2420 | MATa, *ura3*, *leu2*, *his3*, *met15*, [N-terminally FLAG tagged 10% optimal *HIS3*, *URA3*] | This study |
| yJC2421 | MATa, *ura3*, *leu2*, *his3*, *met15*, [N-terminally FLAG tagged 20% optimal *HIS3*, *URA3*] | This study |
| yJC2422 | MATa, *ura3*, *leu2*, *his3*, *met15*, [N-terminally FLAG tagged 30% optimal *HIS3*, *URA3*] | This study |
| yJC2423 | MATa, *ura3*, *leu2*, *his3*, *met15*, [N-terminally FLAG tagged 40% optimal *HIS3*, *URA3*] | This study |
| yJC2424 | MATa, *ura3*, *leu2*, *his3*, *met15*, [N-terminally FLAG tagged 50% optimal *HIS3*, *URA3*] | This study |
| yJC2425 | MATa, *ura3*, *leu2*, *his3*, *met15*, [N-terminally FLAG tagged 60% optimal *HIS3*, *URA3*] | This study |
| yJC2426 | MATa, *ura3*, *leu2*, *his3*, *met15*, [N-terminally FLAG tagged 70% optimal *HIS3*, *URA3*] | This study |
| yJC2427 | MATa, *ura3*, *leu2*, *his3*, *met15*, [N-terminally FLAG tagged 80% optimal *HIS3*, *URA3*] | This study |
| yJC2428 | MATa, *ura3*, *leu2*, *his3*, *met15*, [N-terminally FLAG tagged 90% optimal *HIS3*, *URA3*] | This study |
| yJC2429 | MATa, *ura3*, *leu2*, *his3*, *met15*, [N-terminally FLAG tagged 100% optimal *HIS3*, *URA3*] | This study |
| yJC2488 | MATα, *ura3*, *leu2*, *his3*, *lys2*, [N-terminally FLAG tagged 100% optimal *HIS3*, *URA3*] | This study |
| yJC2489 | MATα, *ura3*, *leu2*, *his3*, *lys2*, [N-terminally FLAG tagged 40% optimal *HIS3*, *URA3*] | This study |
| yJC2490 | MATα, *ura3*, *leu2*, *his3*, *lys2*, [N-terminally FLAG tagged 0% optimal *HIS3*, *URA3*] | This study |
| yJC2491 | MATα, *ura3*, *leu2*, *his3*, *lys2*, [N-terminally FLAG tagged 10% optimal *HIS3*, *URA3*] | This study |
| yJC2492 | MATα, *ura3*, *leu2*, *his3*, *lys2*, [N-terminally FLAG tagged 20% optimal *HIS3*, *URA3*] | This study |
| yJC2493 | MATα, *ura3*, *leu2*, *his3*, *lys2*, [N-terminally FLAG tagged 30% optimal *HIS3*, *URA3*] | This study |
| yJC2494 | MATα, *ura3*, *leu2*, *his3*, *lys2*, [N-terminally FLAG tagged 50% optimal *HIS3*, *URA3*] | This study |
| yJC2495 | MATα, *ura3*, *leu2*, *his3*, *lys2*, [N-terminally FLAG tagged 60% optimal *HIS3*, *URA3*] | This study |
| yJC2496 | MATα, *ura3*, *leu2*, *his3*, *lys2*, [N-terminally FLAG tagged 70% optimal *HIS3*, *URA3*] | This study |
| yJC2497 | MATα, *ura3*, *leu2*, *his3*, *lys2*, [N-terminally FLAG tagged 80% optimal *HIS3*, *URA3*] | This study |
| yJC2498 | MATα, *ura3*, *leu2*, *his3*, *lys2*, [N-terminally FLAG tagged 90% optimal *HIS3*, *URA3*] | This study |

